# Rainbow smelt population responses to species invasions and change in environmental condition

**DOI:** 10.1101/2020.10.08.332205

**Authors:** Rosalie Bruel, J. Ellen Marsden, Bernie Pientka, Nick Staats, Timothy Mihuc, Jason D. Stockwell

**Affiliations:** Rubenstein Ecosystem Science Laboratory, University of Vermont, Burlington, VT, USA; Vermont Fish and Wildlife Department, Essex Junction, VT, USA; US Fish and Wildlife Service, Essex Junction, VT, USA; SUNY Plattsburgh, Plattsburgh, NY, USA

**Keywords:** alewife, demographics, resource competition, habitat scale, predation

## Abstract

Invasive species can have major disruptions on native food webs, yet the impact of species introductions and whether they will become invasive appears to be context-dependent. Rainbow smelt and alewife co-exist as invasive species in the Laurentian Great Lakes and as native species on the Atlantic coast of North America, but in Lake Champlain rainbow smelt is the dominant native forage fish and alewife are invasive. Alewife became abundant by 2007, providing an opportunity to explore the dynamics of these two species in a system where only one is invasive. We used data from a 31-year forage fish survey to compare demographics of rainbow smelt populations in three basins of Lake Champlain with different pelagic volumes, nutrient levels, and predator abundances. Rainbow smelt catch-per-unit-effort (CPUE) remained constant in the large, deep Main Lake before and after alewife invaded, but decreased in the two smaller basins. Declines were primarily a result of increased age-0 and age-1 mortality. Predation by top piscivores, system productivity, and resource competition alone could not explain the patterns in CPUE across the basins. The mechanisms that allow alewife and rainbow smelt to co-exist could be related to system volume and oxythermal habitat availability, and may explain why the two species do not negatively affect each other in other systems. Summer hypoxia in the smaller basins could force individuals into smaller habitat volumes with higher densities of competitors and cannibalistic adult smelt. Our findings suggest that habitat size mediates the impact of invasive alewife on native rainbow smelt.

## Introduction

Effects of invasive species on ecosystems are difficult to predict based on the ecology of the invading species in their native range or by their behavior as an invasive species in other systems (Mackie and Schloesser, 1996). Invasive species may have antagonistic or synergistic effects with other invasive species, although facilitation between invasive species (i.e., the invasion meltdown hypothesis) is most common (Braga et al., 2018; Glon et al., 2017; Simberloff and Von Holle 1999). Native species’ responses to invasive species can range from improved fitness and survival to local extirpation (Jacobs et al., 2017; Madenjian et al., 2008). In aquatic systems, these responses can be related to size of the system, complexity of the existing native community, and disturbance history of the system, including prior loss of native species, habitat degradation, and prior invasions (Brook et al., 2008; Ricciardi and Macisaac, 2010). Comparison of systems in which two species are both native, are both invasive, or one is native and the other invasive, can inform our understanding of the dynamics between native and invasive species.

Rainbow smelt (*Osmerus mordax*) and alewife (*Alosa pseudoharengus*) are native to and coexist along the Atlantic coast of North America (Bigelow and Schroeder, 2002). They also coexist in the Laurentian Great Lakes, where both species invaded in the early 1900s (Mills et al.,1993) and are now important prey for top predators, including lake trout (*Salvelinus namaycush*) and introduced Pacific salmonines (*Oncorhynchus* spp.; Happel et al., 2018; Ray et al., 2007; Stewart et al., 1981). Both species are pelagic planktivores and affect many native species in the Great Lakes because they feed on native larval fishes, compete with other planktivores, and are linked to thiamine deficiency in salmonine predators (Harder et al., 2018; Krueger et al., 1995; Madenjian et al., 2008; Myers et al., 2009). However, the two species do not appear to negatively affect each other, despite spatial overlap of larvae and age-0 life-stages (Madenjian et al., 2008). In some systems, however, alewife are invasive and rainbow smelt are native, and such a context provides the opportunity to further examine their interactions. Here, we focus on the effects of a recent invasion of alewife on a native rainbow smelt population in Lake Champlain, USA.

Lake Champlain has a relatively intact biotic community, with only two species extinctions (lake trout and Atlantic salmon, but reintroduced by stocking), and a small number of introduced species (51) relative to the Great Lakes (at least 188; Lake Champlain Basin Program, 2018; Marsden and Hauser, 2010; Marsden and Langdon, 2012; Ricciardi, 2006). The native coldwater prey fish community in Lake Champlain has low diversity, consisting primarily of rainbow smelt, trout-perch (*Percopsis omiscomaycus*), slimy sculpin (*Cottus cognatus*), and cisco (*Coregonus artedi*). The addition of alewife as an alternative prey could release rainbow smelt from predation and indirectly lead to increased rainbow smelt abundance. However, predatory release might eventually lead to higher rainbow smelt cannibalism, as 38-93% of age-0 mortality in the lake prior to alewife invasion has been attributed to cannibalism (Parker Stetter et al., 2007).

Rainbow smelt in Lake Champlain supported native populations of landlocked Atlantic salmon (*Salmo salar*), lake trout and walleye (*Sander vitreus*) (Marsden and Langdon, 2012). Unlike the Great Lakes, where rainbow smelt spawn in tributaries, spawning in Lake Champlain occurs offshore at depths greater than 15 m. Since the extirpation of the two salmonine species in the 1800s and the decline of walleye in the 1900s, rainbow smelt populations appeared to be regulated by cannibalism and intra-specific competition rather than predation by stocked salmonines (He and Labar, 1994; Kirn and Labar, 1996; Labar, 1993; Parker-Stetter et al., 2007; Stritzel-Thomson et al., 2011), in spite of sustained stocking of lake trout and Atlantic salmon that began in 1973. After two decades of high stocking levels, lake trout stocking has been maintained at an average of 83,400 yearling equivalents since 1996 (Fisheries Technical Committee, 2016, 2008). Larger numbers of Atlantic salmon are stocked annually (an average of 278,000 yearling equivalents since 1987). Steelhead trout (*Oncorhynchus mykiss*) and brown trout (*Salmo trutta*) are also stocked at numbers similar to lake trout (Pientka unpublished data). Walleye stocking in three tributaries started in the late 1980s.

Alewife were first discovered in Lake Champlain in 2003 and became abundant by 2007 (Marsden and Hauser, 2009), and have been incorporated into diets of walleye, Atlantic salmon, and to a lesser extent, lake trout (Simonin et al., 2018). Since the alewife invasion, body size of two zooplankton groups decreased to or below the size of alewife feeding preference (Mihuc et al., 2012), suggesting alewife could indirectly suppress rainbow smelt through competition for zooplankton (Kircheis et al., 2004, Urban and Brant 1993). Adult alewife could also directly suppress rainbow smelt through predation on larvae as the two life stages spatially overlap during summer (Simonin et al., 2012).

The general understanding of alewife invasions in large lakes is based on the Great Lakes, where both alewife and rainbow smelt replaced overfished coregonine planktivores and do not appear to impact each other (Madenjian et al., 2008). In Lake Champlain, the presence of a relatively intact fish community with a robust population of rainbow smelt suggests the forage fish community could be resilient to additional invasion. Alternatively, relatively low prey fish diversity in Lake Champlain could make the community vulnerable to an alewife invasion; therefore we hypothesized that alewife would negatively impact rainbow smelt. We tested our hypothesis using long-term survey data from three separate and semi-isolated basins of Lake Champlain (the Main Lake, Malletts Bay, and Northeast Arm) which differed in predator composition and abundance, productivity, size, and oxythermal conditions (Table 1). Rainbow smelt populations in the three basins have different demographic structures that indicate the populations are largely isolated from each other, although they are not genetically different (Euclide et al., 2020). Specifically, we predicted that, after alewife invaded,: (1) based on predator abundance, rainbow smelt abundance measured as catch-per-unit-effort (CPUE) would decrease more in the Main Lake than the two smaller basins where lake trout are absent for much of the year; (2) based on productivity, rainbow smelt CPUE should have the smallest change in the Northeast Arm, because higher food availability would reduce competition; and (3) based on basin size, rainbow smelt CPUE should decrease the most in Malletts Bay, because small volume increases spatial overlap (Evans and Loftus 1987; Latta 1995), oxythermal habitat may be limited (Hrycik et al., 2017), and alewife:rainbow smelt interactions may include competition or predation. Interspecific predation largely occurs during the larval stage (Simonin et al., 2019); thus (4) we expected that if alewife affect only age-0 rainbow smelt, then rainbow smelt mortality rate past age-1 would not change. Finally, (5) if competition between rainbow smelt and alewife was high and rainbow smelt was released from predation, we would expect rainbow smelt condition and length to decline in basins where alewife became abundant. Overall, our objective was to determine whether and how invasive alewife affected native rainbow smelt in a relatively intact large-lake community.

**Table 1.** Characteristics of the three major basins in Lake Champlain. Basin morphometry data from Myer and Gruendling (1979); total phosphorus (TP) data from Smeltzer et al., (2012); predator abundance data from Bernie Pientka (unpublished data).

|  | <u>Main Lake</u> | <u>Malletts Bay</u> | <u>Northeast Arm</u> |
| --- | --- | --- | --- |
| Predator abundance | high | medium | medium |
| Productivity ($\mu\text{g/l TP}$) | low (10-15) | low (8-12) | high (20-25) |
| Basin volume ($\text{km}^3$) | large (21.0) | small (0.72) | medium (3.45) |
| Max. basin depth (m) | 122 | 32 | 49 |
| Avg basin depth (m) | 30.8 | 13.3 | 12.8 |

## Material and methods

### Study system

Lake Champlain is a large lake (26 km^3^ and 1,130 km^2^) located among Vermont, New York (US), and Québec (CAN; Fig. 1). The Main Lake extends from Crown Point (NY) at the south to Rouses Point (NY) at the north, and contains the largest volume and the deepest areas of the lake (Table 1). Malletts Bay and the Northeast Arm are isolated from each other and the Main Lake by large islands and several causeways up to 5.2 km long between islands and the mainland. Water exchange and fish passage are possible through shallow, narrow connections in each causeway (see Fig. 3 in Marsden and Langdon, 2012). The three basins differ in their nutrient levels and total volume (Table 1). The Northeast Arm is the most productive basin and has an extensive hypoxic zone that limits available summer habitat for lake trout. The Main Lake is the largest basin and has moderate to low productivity, and Malletts Bay is the smallest and least productive basin. Major fish predators in all three basins include Atlantic salmon, burbot (*Lota lota*), and walleye. Lake trout are present in all basins in winter but not in Malletts Bay and the Northeast Arm during summer (Pientka, unpublished data), as temperatures in the causeway passages exceed their thermal optima and oxythermal habitat in the smaller basins is limited in summer.

**Figure 1.**
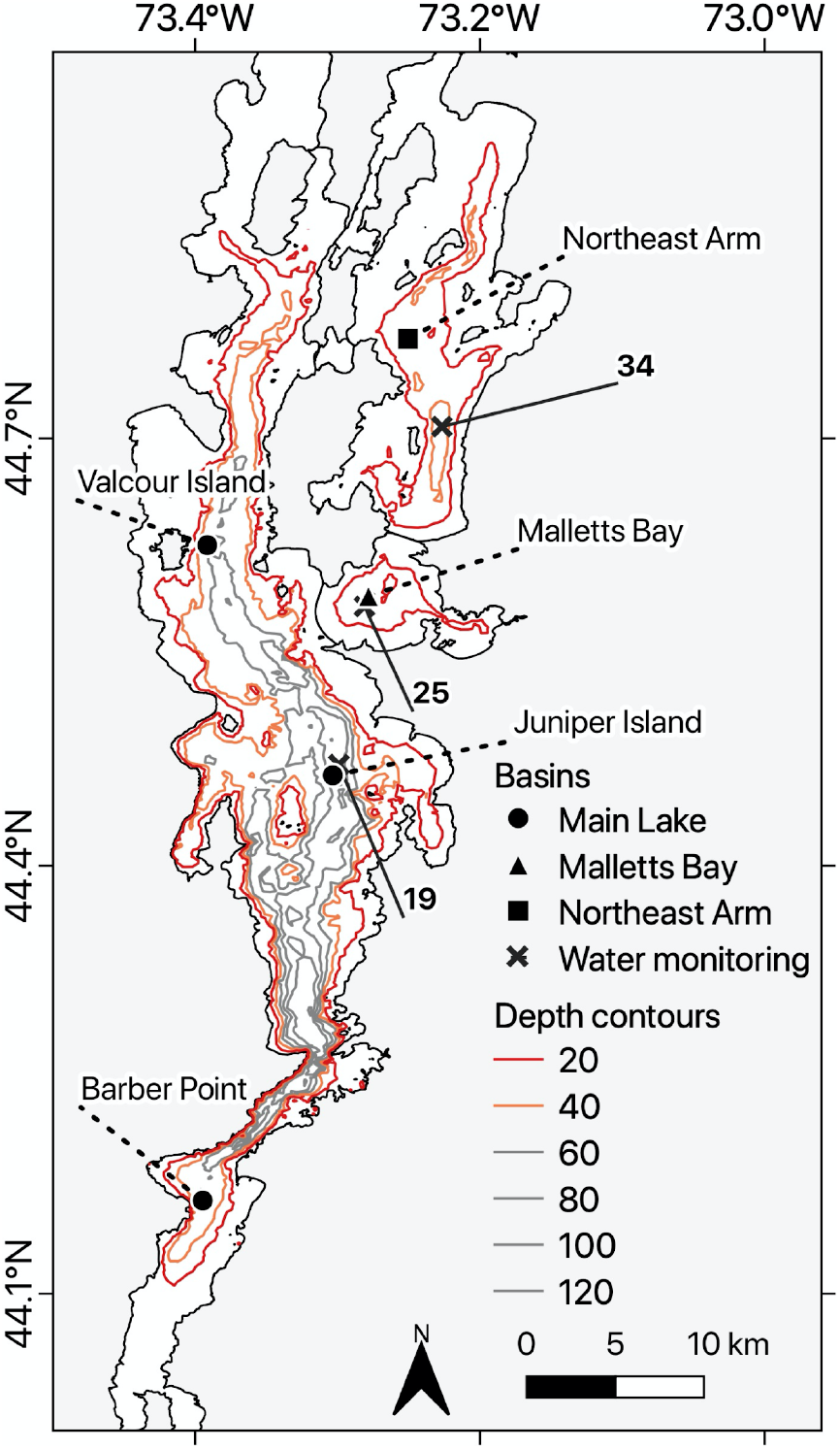
Lake Champlain bathymetry and basins, with trawling and long-term monitoring stations.

### Long-term survey data

The Vermont Fish and Wildlife Department conducted standardized assessment sampling of rainbow smelt from 1990 to 2015. The five standard stations included one in each of Malletts Bay and the Northeast Arm, and three in the Main Lake: north (Valcour Island), central (Juniper Island), and south (Barber Point; Figure 1). The most consistent sampling was conducted using stepped-oblique midwater trawling at night from late July through mid-August, when young-of-year (YOY) and yearling-and-older (YAO) fishes were vertically separated by thermal stratification (Table 1; Labar, 1998). Trawls began at 35-m depth or just above lake bottom (26 m in Malletts Bay and 29 m in the Northeast Arm) and were fished for 10 minutes, raised approximately 3 m and fished again for 5 min at that depth, continuing in a steplike fashion until the net reached 10 m below the surface (Labar, 1998). Four trawls were conducted at each of the five standard stations. CPUE was calculated as catch per 55 min of trawling. Fifty rainbow smelt were randomly sampled from each trawl and frozen on board, so up to 200 individuals were used each year per station to collect population demographics. In the laboratory, rainbow smelt were measured for total length (TL) and weight. Otoliths were extracted and stored in an ethanol/glycerol mixture (70:30) and age was estimated by counting annuli using whole otoliths under 10-45x magnification. Floating gillnets were added in 2008 to focus on capture of YOY and YAO alewife which are undersampled in the midwater trawl. Nets were set at standard stations before dark and fished for four hours, and CPUE was calculated as catch per four hours.

We used vertical profile data from the Lake Champlain Long-Term Water Quality and Biological Monitoring Project to describe the oxythermal habitat in each basin (https://dec.vermont.gov/watershed/lakes-ponds/monitor/lake-champlain). We selected three stations, one in each basin, sampled fortnightly from late April to early November, to represent conditions in the Main Lake, Malletts Bay, and the Northeast Arm (Fig. 1). We also analyzed changes in mean summer (July-August) zooplankton density collected by the Lake Champlain long-term monitoring program since 1992 at the same stations. Zooplankton samples were collected with whole water vertical tows taken monthly or bi-weekly using a 30-cm diameter, 153-um mesh net during the day (Mihuc et al., 2012). Zooplankton were identified to the lowest possible taxon. For the most abundant taxa (abundance > 5%) at the Main Lake and Northeast Arm stations, 7-10 individuals were measured per sample to estimate average length each year since 2001.

### Data analysis

Annual data for rainbow smelt were pooled by basin and the resulting means were used for all the analyses. We tested for differences in CPUE among periods: 1987-2002 (before alewife invasion), 2003-2006 (invasion), and 2007-2015 (after invasion). We also tested for differences in average length (age-2+, because ages 0 and 1 were not well recruited into the gear) and Fulton’s condition factor among periods for each basin using a Kruskal-Wallis pairwise comparison test. Fulton’s condition, calculated for each individual, is the weight of a given individual divided by the cube of its length (Ricker, 1975). A scaling factor of 10^−6^ was applied to bring the condition close to 1. We used age-2 and age-3 fish for this analysis because condition cannot be compared across too many age classes due to allometric growth (Guy and Brown, 2007). We did not include weight comparisons because average length and weight were highly correlated (Pearson’s product-moment correlation = 0.94).

We used longitudinal data (ages 2-5) of each cohort to calculate annual mortality rate (A). We estimated year-class CPUE in each year using the proportion of each year class in the annual subsamples of 200 individuals at each station, multiplied by the CPUE of that station that year. We calculated the instantaneous mortality rate Z as the slope of the linear relationship between age and the natural log of CPUE (Ogle et al., 2020):

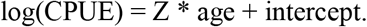

We excluded age-0 and −1 because these ages were not fully recruited to the gear, and age-6 and older because none were collected after the alewife invasion. A is calculated as:

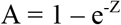

(Ogle et al., 2020). We compared annual mortality between the periods 1995-2002, before the arrival of alewife, and 2007-2015, after alewife became abundant. We used a Kruskal-Wallis pairwise comparison test for each of the three basins separately, and applied a Bonferroni correction of α/3 for significance.

We calculated the maximum habitat size for rainbow smelt using a sub-lethal threshold of 4.5 mg/l of dissolved oxygen (DO; Hrycik et al., 2017), and a temperature threshold of 16°C (Lantry and Stewart, 1993). For each sampling date, we extracted the shallowest depth for which temperature was below 16°C, and the deepest depth with DO concentration above 4.5 mg/l. The difference between these depths was used as an estimate of habitat availability for rainbow smelt.

Average densities of abundant zooplankton taxa each year were averaged before (1992-2002), during (2003-2006) and after (2007-2015) alewife invasion. Similarly, average length was estimated for 2001-2002, 2003-2006, and 2007-2015. For both zooplankton densities and lengths, we tested for differences among the three time periods using a Kruskal-Wallis pairwise comparison test.

All computational work and visualization was done using the packages *FSA* (Ogle et al., 2020), *ggplot2* (Wickham, 2016), and *PerformanceAnalytics* (Peterson and Carl, 2020), with R version 3.6.3 (R Core Team, 2020).

## Results

Rainbow smelt CPUE was, on average, highest in the Northeast Arm (mean ± SD = 967 ± 1,095 fish per 55-min trawl), lowest in the Main Lake (265 ± 320), and intermediate in Malletts Bay (772 ±1,103) between 1987 and 2015 (Fig. 2A, Table 2). CPUE in Malletts Bay and the Northeast Arm, however, were both lower in 2007-2015 than prior to and during alewife invasion (p < 0.01 for Malletts Bay and p < 0.05 for the Northeast Arm, Fig. 2A, Supplementary Material 1). CPUE increased significantly in Malletts Bay until 2002 (slope = 0.1, p = 0.004) and remained stable in the Northeast Arm (slope = 0.04, p = 0.59) until 2002, and decreased after 2003 (p < 0.001 in both basins, Figure 2B). CPUE remained unchanged in the Main Lake over the same time period (Figure 2). Overall, rainbow smelt CPUE declined 100-fold in the Northeast Arm and 30-fold in Malletts Bay after alewife became established in 2007. Alewife catches in floating gillnets were heterogeneous and the data should be treated with caution. However, the data support the field observation that alewife were consistently present in all basins (Fig. 3).

**Figure 2.**
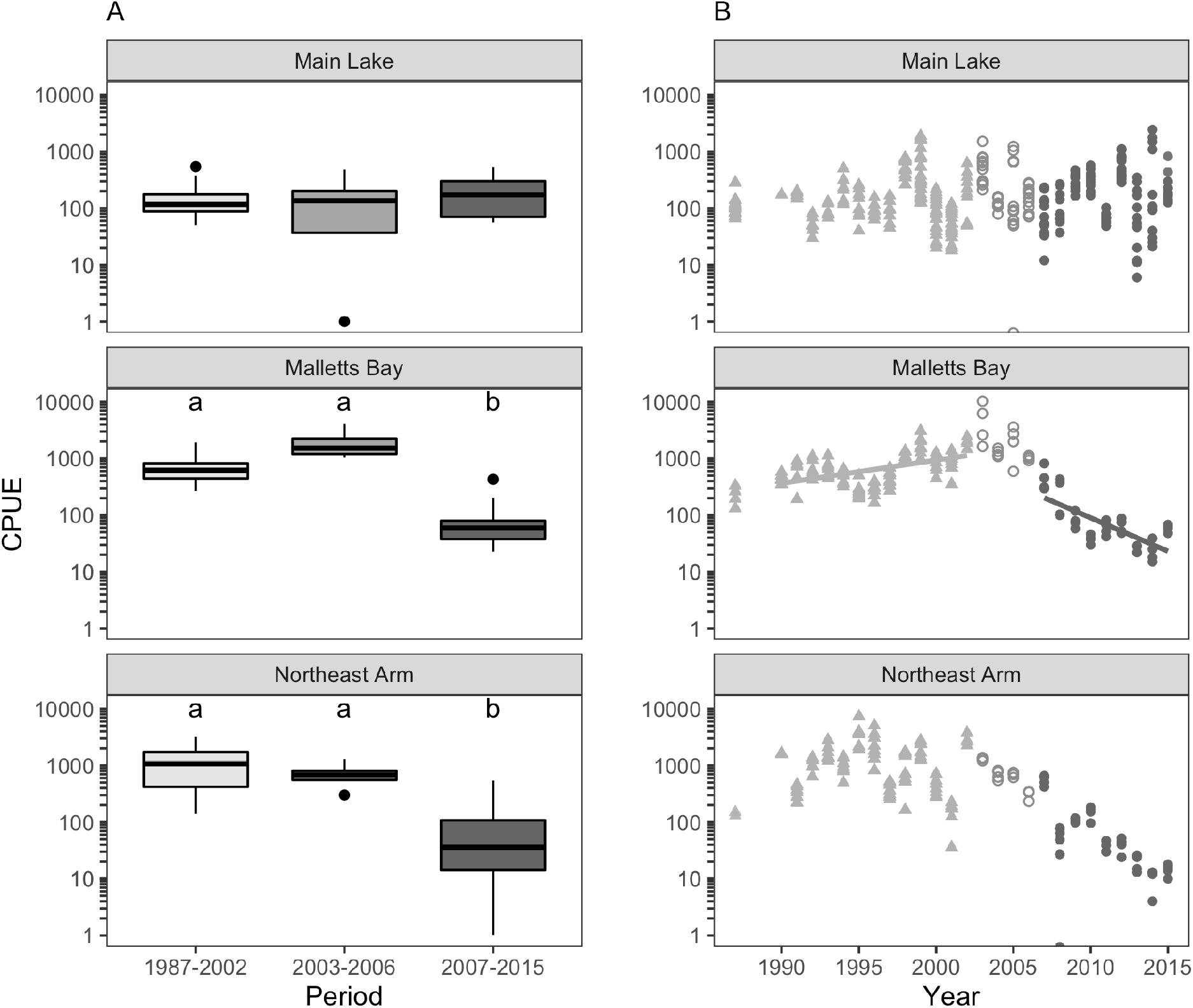
**(A)** Boxplot of average rainbow smelt CPUE and **(B)** changes in CPUE by period in each basin. Limits of each box represent the 25-75% quantiles, dark bars represent the median, lines show the 10-90 % limits of the CPUE, and dots represent outliers from the 10-90% distribution. The y-axis scale is logarithmic. Letters indicate groups that are significantly different (Kruskal-Wallis test with Bonferroni correction).

**Table 2.** Characteristics of sampling station (maximum depth, typical starting depth for trawling, and trawling time) in Lake Champlain with average catches of rainbow smelt standardized to 55 min trawl time at each station each year. Number in parentheses is the number individuals kept for biological analyses. A ‘-’ indicates data or subsamples were not collected. No sampling was conducted in 1988-1989.

| Basin | Main Lake |  |  | Malletts Bay | Northeast Arm |
| --- | --- | --- | --- | --- | --- |
| Station | <u>Barber Point</u> | <u>Juniper Island</u> | <u>Valcour Island</u> | <u>Malletts Bay</u> | <u>Northeast Arm</u> |
| max depth (m) | 50-60 | 70-90 | 56-62 | 22-32 | 22-40 |
| starting depth (m) | 35 | 35 | 35 | 26 | 29 |
| trawling time (min) | 55 | 55 | 55 | 40 | 45 |
| Year | Mean CPUE |  |  |  |  |
| 1987 | 139 (-) | 110 (-) | - | 231 (-) | 139 (-) |
| 1990 | - | 175 (119) | - | 448 (199) | 1,628 (125) |
| 1991 | - | 173 (172) | - | 614 (125) | 324 (191) |
| 1992 | - | 53 (176) | - | 654 (199) | 1,104 (192) |
| 1993 | 93 (49) | - | - | 654 (201) | 1,674 (205) |
| 1994 | 316 (284) | 126 (101) | - | 461 (203) | 977 (100) |
| 1995 | 202 (92) | 72 (93) | - | 278 (195) | 3,553 (217) |
| 1996 | 79 (100) | 111 (100) | - | 305 (200) | 2,440 (200) |
| 1997 | 124 (57) | 66 (-) | - | 465 (202) | 398 (199) |
| 1998 | - | 572 (303) | - | 1,127 (196) | 1,069 (208) |
| 1999 | 317 (200) | 146 (200) | 1,288 (200) | 1,696 (200) | 1,814 (200) |
| 2000 | 65 (175) | 55 (199) | 155 (198) | 864 (200) | 440 (195) |
| 2001 | 63 (100) | 84 (201) | 35 (138) | 864 (198) | 158 (183) |
| 2002 | 247 (203) | 51 (168) | 374 (200) | 1,957 (200) | 2,869 (196) |
| 2003 | 587 (200) | 825 (200) | 285 (200) | 5,193 (200) | 1,304 (200) |
| 2004 | 138 (203) | 113 (200) | - | 1,276 (200) | 690 (200) |
| 2005 | 902 (200) | 50 (194) | 78 (198) | 2,226 (199) | 693 (200) |
| 2006 | 256 (201) | 131 (196) | 121 (199) | 1,037 (200) | 306 (179) |
| 2007 | 152 (162) | 77 (172) | 56 (210) | 470 (200) | 551go (200) |
| 2008 | 209 (202) | 67 (194) | 64 (210) | 252 (224) | 49 (201) |
| 2009 | 300 (206) | 400 (197) | 248 (198) | 82 (224) | 108 (199) |
| 2010 | 459 (199) | 305 (200) | 185 (201) | 38 (104) | 150 (202) |
| 2011 | 66 (199) | 67 (214) | 71 (187) | 61 (164) | 36 (114) |
| 2012 | 811 (205) | 388 (201) | 595 (205) | 66 (235) | 40 (155) |
| 2013 | 12 (48) | 254 (197) | 68 (200) | 25 (69) | 20 (63) |
| 2014 | 1,697 (193) | 117 (160) | 29 (115) | 24 (67) | 10 (32) |
| 2015 | 630 (200) | 223 (200) | 153 (199) | 60 (162) | 14 (45) |

**Figure 3.**
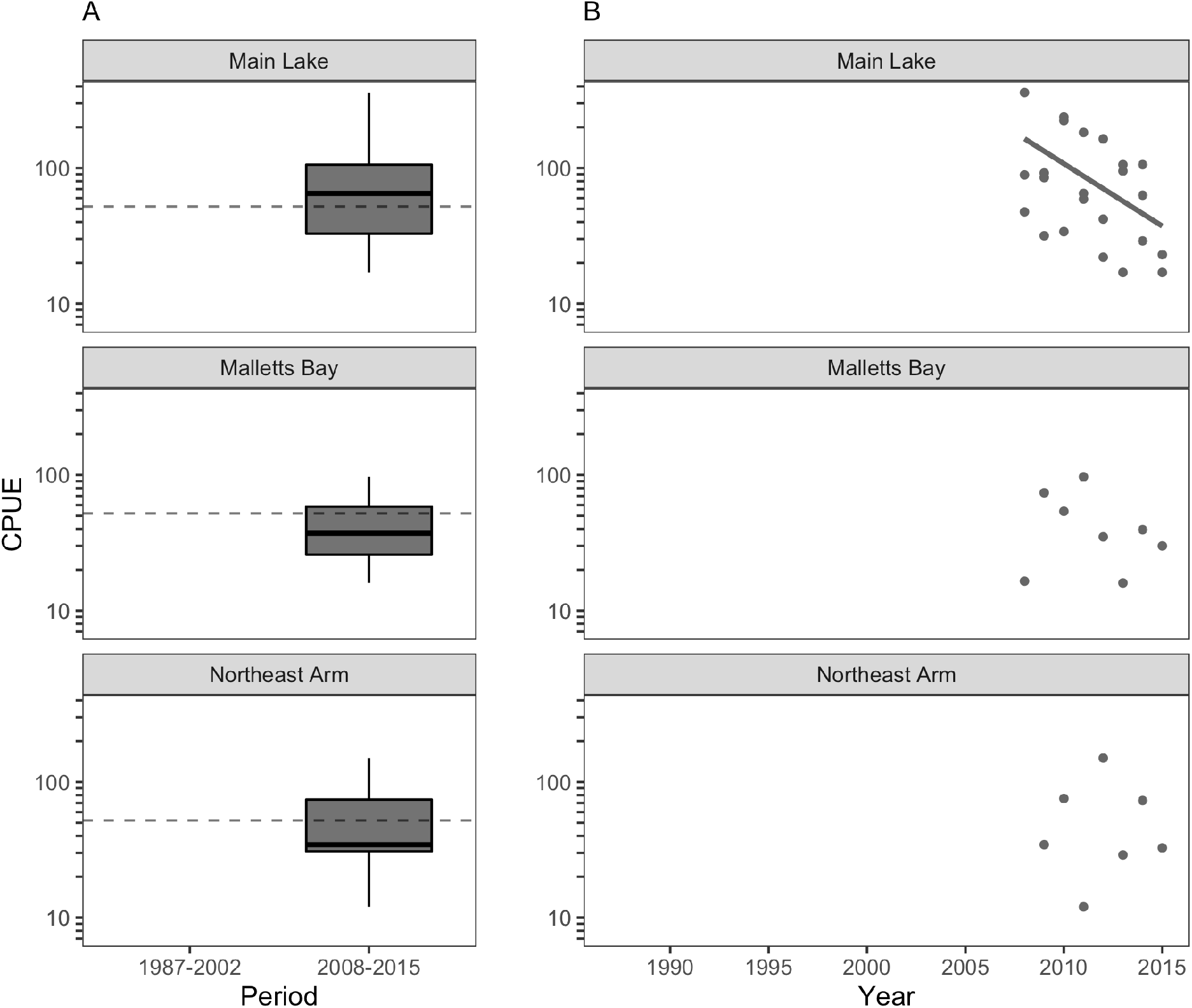
**(A)** Boxplot of average alewife CPUE and **(B)** changes in CPUE in each basin. See Fig. 2 for description of boxplot. Horizontal dotted line represents average CPUE. YOY and YAO caught by floating gillnets were summed together.

Average length of age-2+ rainbow smelt was not different between periods except in the Northeast Arm, where length increased between 1987-2002 and 2007-2015 (p < 0.0001, Fig. 3A, Supplementary Materials 1). Average length in 1987-2002 was highest in the Main Lake (147 ± 11.1 mm), lowest in Malletts Bay (127 ± 7.5 mm) and intermediate in the Northeast Arm (134 ± 9.1). In 2007-2012, after alewife became abundant, average length remained overall lowest in Malletts Bay (134 ± 14.2 mm), intermediate in the Main Lake (138 ± 13.8 mm) and highest in the Northeast Arm (153 ± 15.1 mm). Variability in length increased in all basins after alewife became established (Fig. 4). Condition was not significantly different among basins in each period (Fig. 4B). Annual mortality was higher in only one comparison, between 1995-2002 and 2003-2006 in Malletts Bay (p = 0.007, Fig. 4C, Supplementary Material 1).

**Figure 4.**
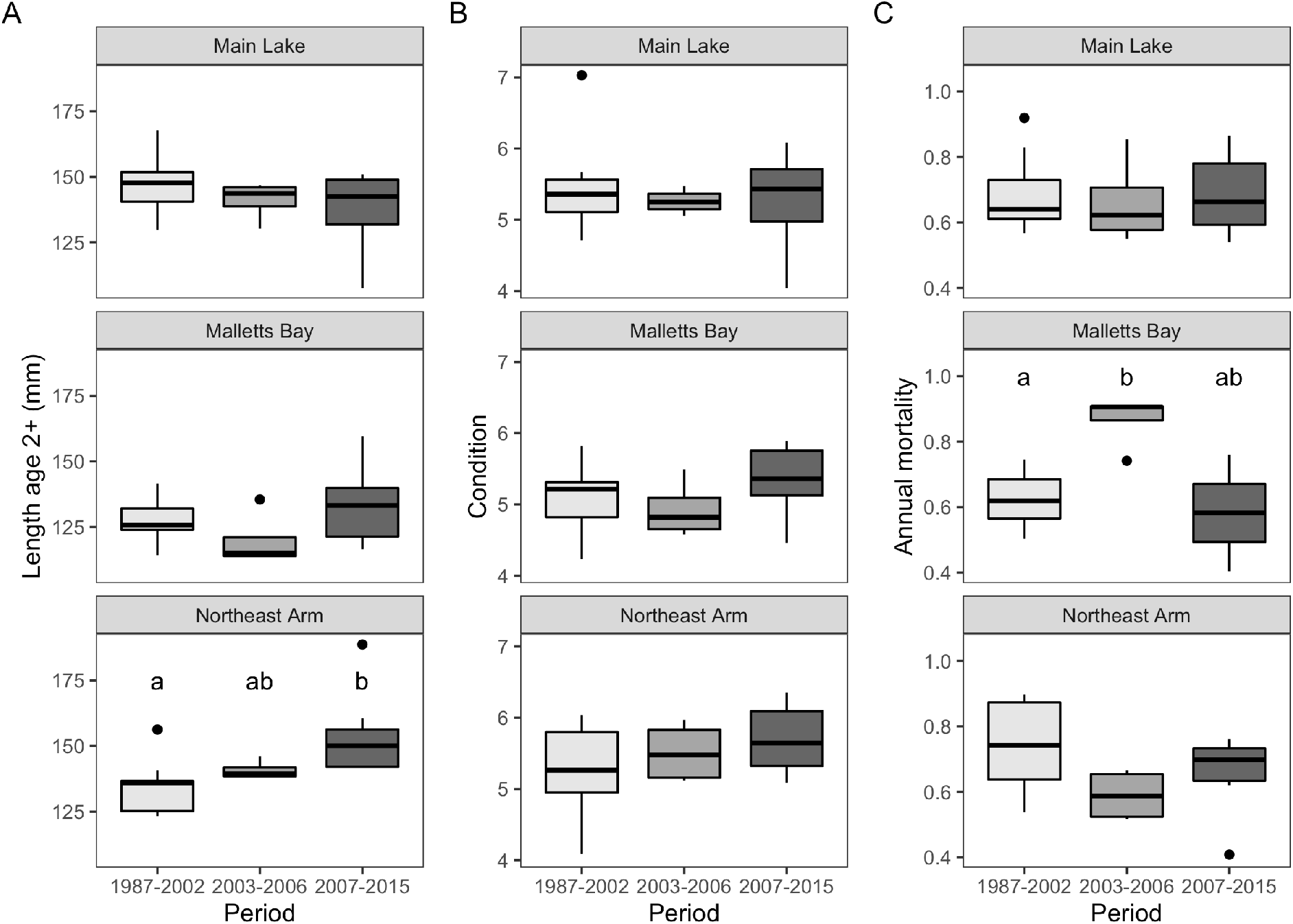
**(A)** Average total length of rainbow smelt (age-2+) in three basins of Lake Champlain for three survey periods, **(B)** average condition for age 2 and 3 rainbow smelt, and **(C)** annual mortality per cohort. Letters indicate groups that are significantly different (Kruskal-Wallis test with Bonferroni correction).

The three basins stratify during the summer, with available oxythermal habitat more restricted in Malletts Bay and the Northeast Arm than in the Main Lake (Fig. 5, Supplementary Material 2). DO concentration remained above 5 mg/l between April and November at all depths in the Main Lake, but was below 4.5 mg/l every summer in Malletts Bay and the Northeast Arm (Fig. 5, Supplementary Material 2). The depth of the epilimnion increased during the summer in every basin, limiting the available near-surface habitat. Overall, conditions were more stressful for rainbow smelt in the 1992-2002 period, during which the habitat was restricted to 4 m or less 73% of the days that were sampled between August and October in the Northeast Arm, and 52% of the time in Malletts Bay. The period of unfavorable conditions dropped to 36 and 41% in the Northeast Arm and Malletts Bay respectively during the 2007-2015 period (Fig. 6).

**Figure 5.**
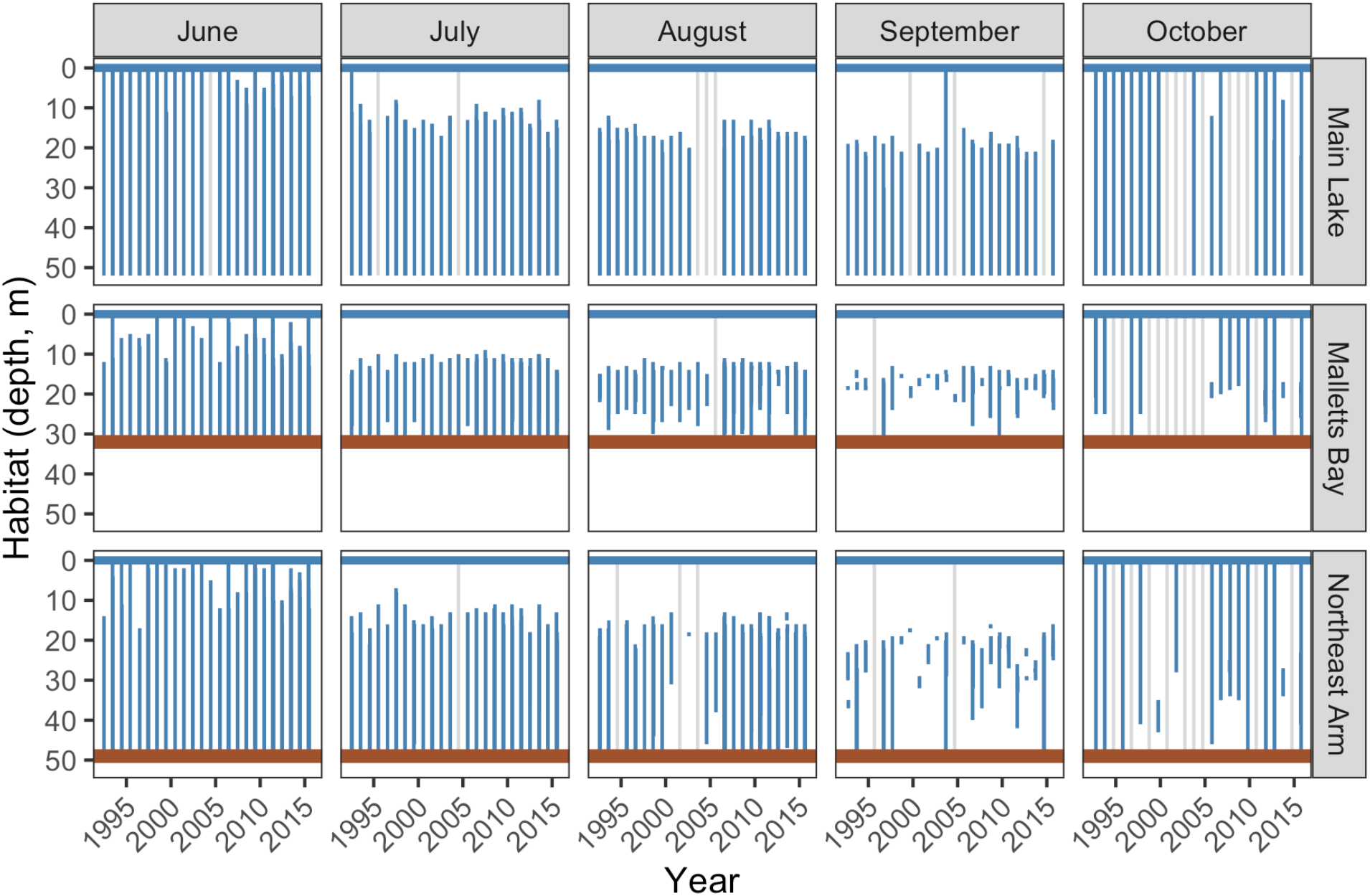
Habitat available for rainbow smelt per month, basin, and year, expressed by depths with oxygen > 4.5 mg/l and temperature < 16°C. Horizontal lines represent lake surface and bottom sediment (not visible for the Main Lake station because depth is 102 m). Lighter grey vertical lines indicate absence of data.

**Figure 6.**
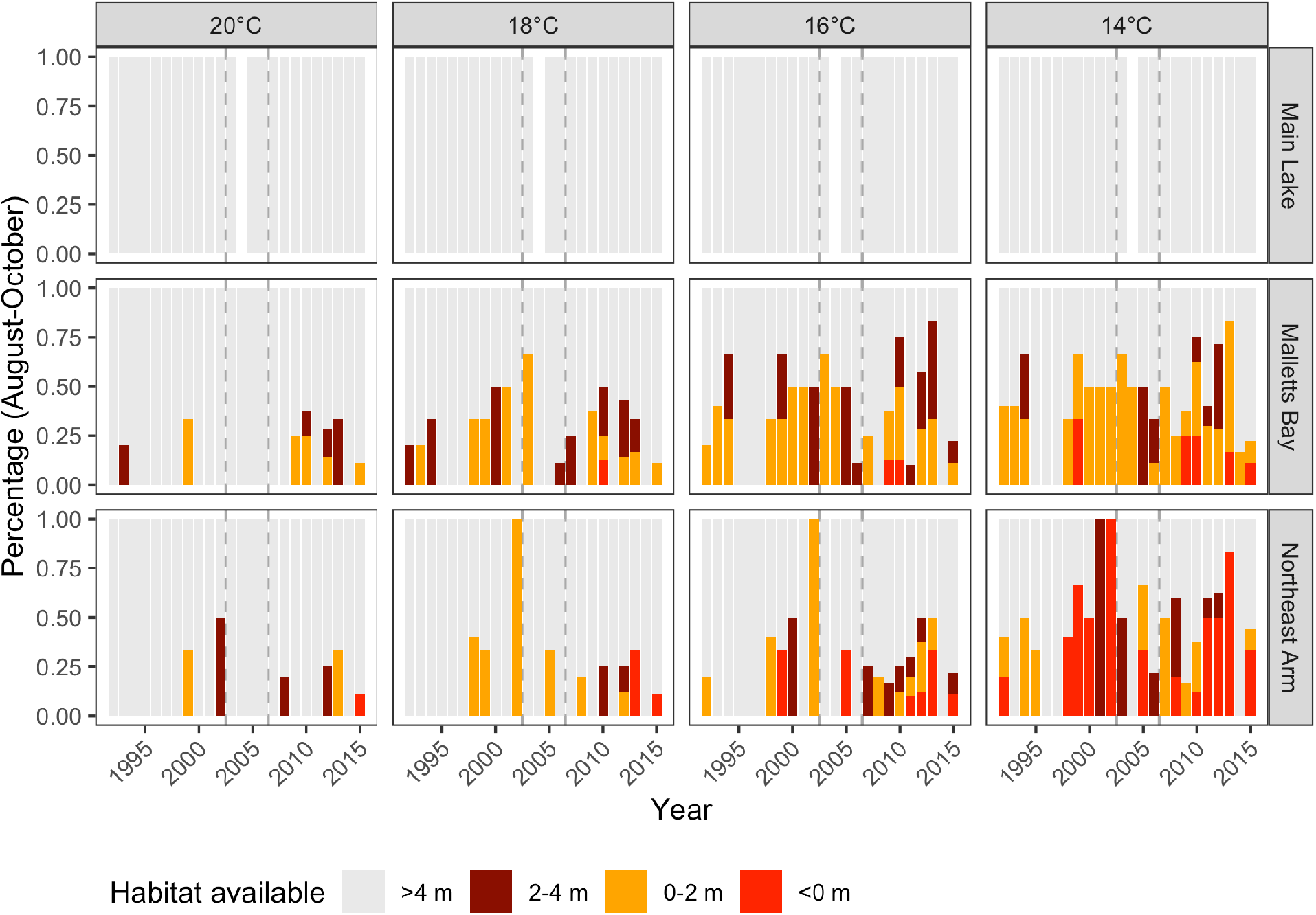
Percentage of days in August-October per year and per basins for which suitable water column habitat (defined by temperature below four possible thresholds and > 4.5 mg O2/L) was between 2-4 m, 0-2 m, or unavailable (0 m). Dashed vertical lines indicate the separation between the periods used for Fig. 2-4.

Zooplankton densities remained stable across the study periods at all three stations with the exception of declines in calanoid copepods and *Daphnia* sp. during the alewife colonization period (2003-2006) (Figure 7). Zooplankton body size did not change before and after the alewife invasion in the Main Lake and Northeast Arm, with the exception of *Daphnia retrocurva* which exhibited a decrease in body size after 2006 in the Main Lake (average length per period: 2001-2002 = 0.96 ± 0.04 mm, 2003-2006 = 1.03 ± 0.15 mm, 2007-2015 = 0.77 ± 0.11 mm; difference in means was between 2003-2006 and 2007-2015, p = 0.050; Figure 8).

**Figure 7.**
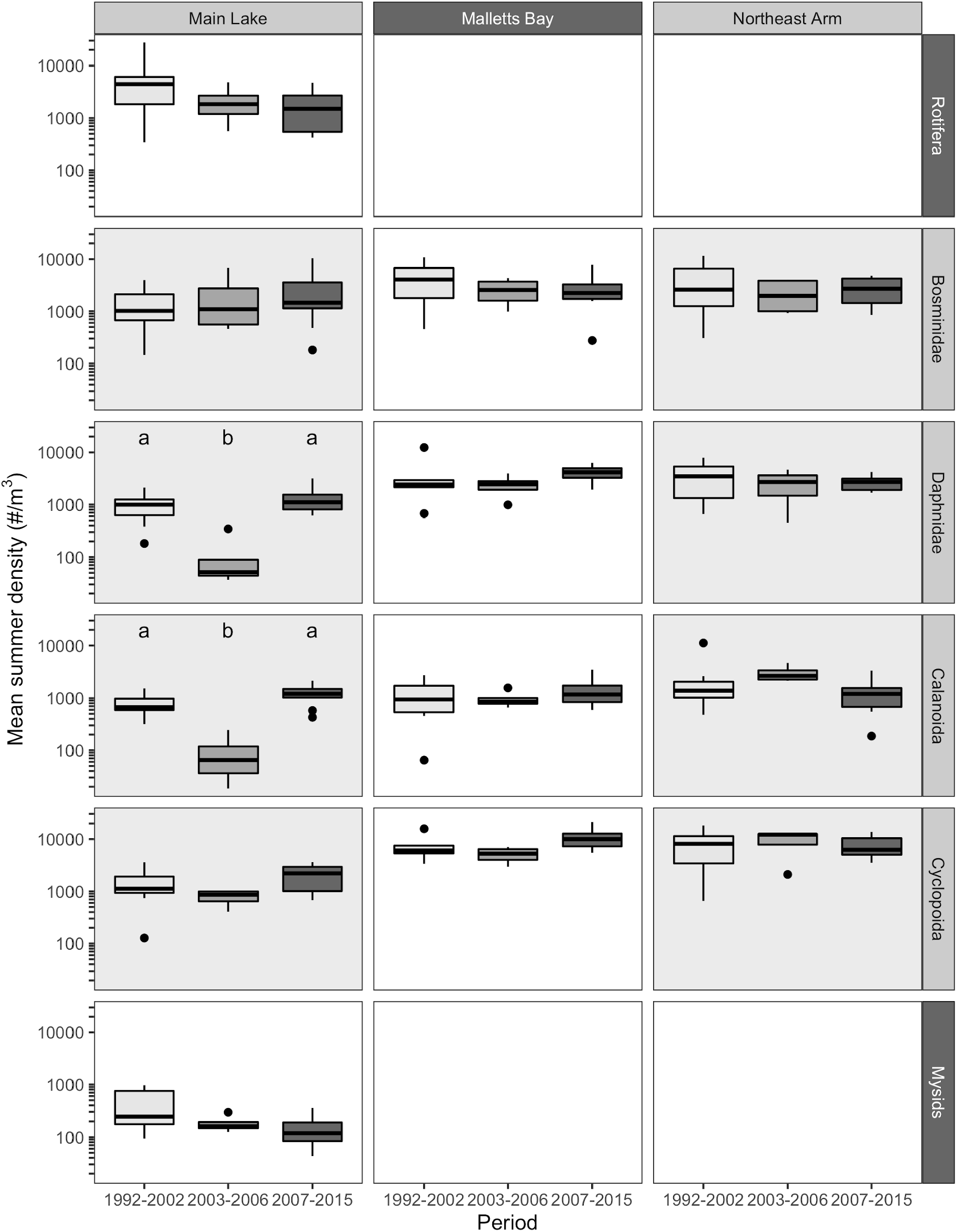
Mean summer density (#/m^3^) of most abundant zooplankton groups, per basins. Grey background in plots indicate available length data (see Fig. 8).

**Figure 8.**
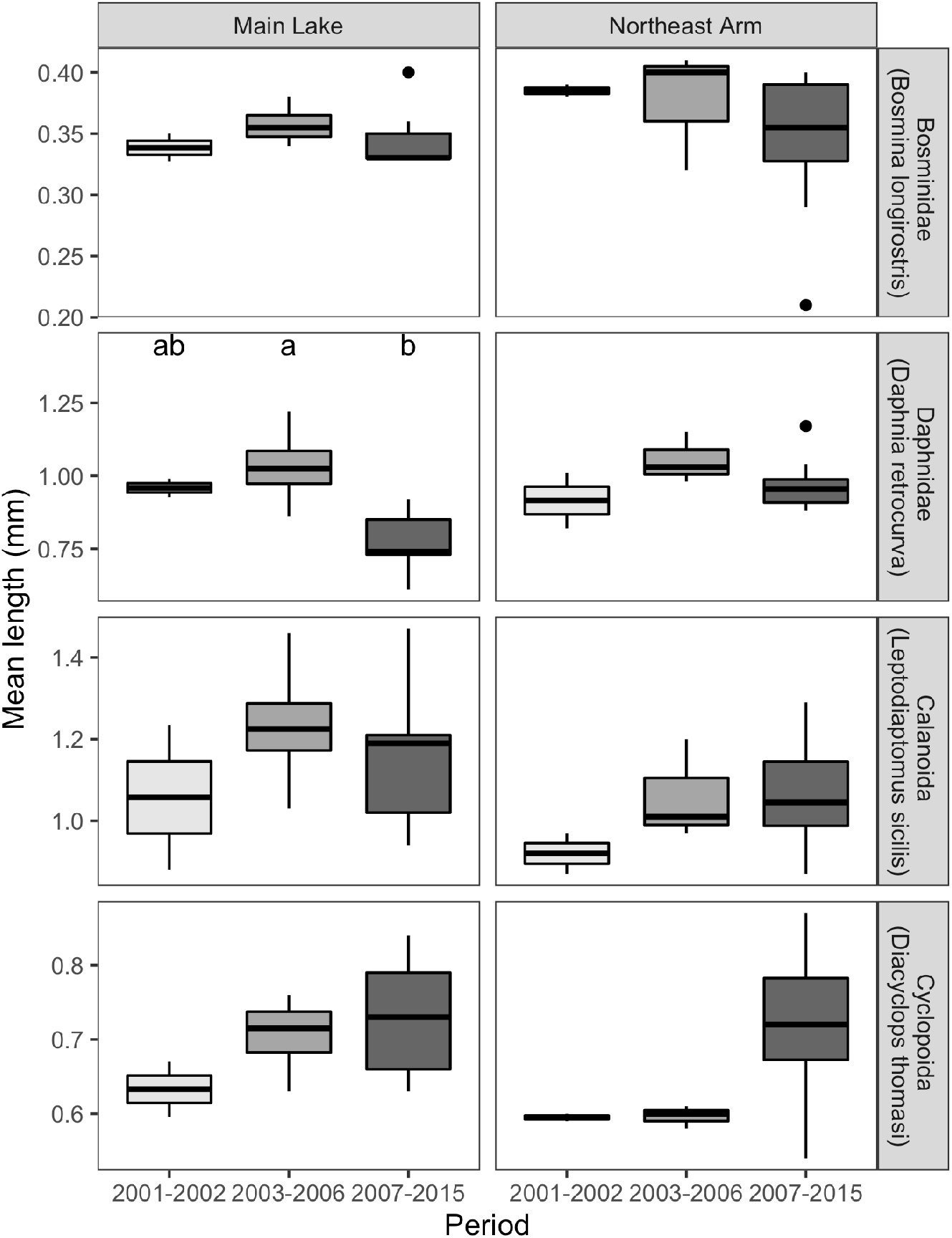
Mean length (mm) of most abundant zooplankton taxa, per basin.

## Discussion

Our basic hypothesis that the alewife invasion of Lake Champlain would negatively impact rainbow smelt was supported in two of our three study basins, in contrast with expectations drawn from the Great Lakes (Madenjian et al., 2008). Rainbow smelt CPUE declined sharply in Malletts Bay and the Northeast Arm, but remained stable in the Main Lake basin after alewife became established. Our prediction that the absence of lake trout in the two smaller basins would result in lower declines than in the Main Lake was not supported. We also predicted that the high productivity in the Northeast Arm would support higher rainbow smelt abundance even after the addition of competition with alewife, but the decline in rainbow smelt CPUE was as severe in the Northeast Arm as in the less productive Malletts Bay. Adult mortality remained constant before and after alewife invasion, suggesting that the changes in CPUE were due to mortality at age-0 and age-1, likely a consequence of predation, cannibalism, or competition. Absence of a change in average length or condition before and after the alewife invasion suggests that competition remained constant as rainbow smelt CPUE declined. The CPUE decline in the two smaller basins, compared to stable rainbow smelt CPUE in the larger Main Lake, suggests that habitat size may exacerbate the impacts of alewife.

Predation by piscivores does not explain the patterns we observed. Predator abundance would need to have remained stable in the Main Lake and increased in the Northeast Arm. In fact, predator abundance in the Main Lake decreased in the pre-alewife period, when lake trout stocking was reduced from an average of 185,900 to 83,400 equivalent yearlings in 1996, while Atlantic salmon stocking remained at 278,000 yearling equivalents annually. Walleye declined steadily since 1960 (Marsden and Langdon 2012) and stocking was initiated in the late 1980s. Of the three stocked tributaries, one is in the Main Lake and the others are in the South Lake and Missisquoi Bay (north of the Northeast Arm). Walleye survival has not been evaluated, but survival of salmonines increased as a consequence of sea lamprey suppression in the mid-1990s (Marsden et al., 2003). Yet during 1987-2002, rainbow smelt CPUE remained relatively constant or increased in our three study basins, and only began to decline in the Northeast Arm and Malletts Bay in 2007, where lake trout are absent during the stratified period. Predation by adult alewife on young rainbow smelt is another possible mechanism to explain the apparent decline in age-0 and age-1 rainbow smelt in the Northeast Arm and Malletts Bay. A predation model based on seasonal vertical distributions of alewife and rainbow smelt, YOY growth rates, and gape-limitation of adult alewife predicted higher mortality of YOY rainbow smelt in the presence of invasive alewife in Lake Champlain (Simonin et al., 2019). However, predation by adult alewife as a driving force of rainbow smelt dynamics remains to be tested, as we found no evidence in the published literature that alewife consume YOY rainbow smelt (e.g., Brandt, 1980; Stewart and Binkowski, 1986 and references therein; Stewart et al., 2009).

If bottom-up effects of system productivity were important to mitigate any possible impacts of competition from alewife (Power, 1992), we should have observed the most impact on rainbow smelt in the least productive basin, Malletts Bay, and the least impact in the most productive basin, the Northeast Arm. However, the patterns were inconsistent or reversed from these expectations; rainbow smelt CPUE declined and average length and condition did not change before and after alewife invasion in these two basins. In fact, CPUE was lowest in the highly productive Northeast Arm after the invasion. The lower abundance of rainbow smelt and higher productivity in the Northeast Arm should have reduced intra-and inter-specific competition and increased growth and condition of rainbow smelt, compared to Malletts Bay, but we did not observe such patterns.

Alewife and rainbow smelt can be intense competitors for zooplankton (Evans and Loftus, 1987). System size, however, may mitigate or exacerbate competition. In small lake systems where habitat availability and heterogeneity may be limited, alewife appear to outcompete native rainbow smelt (Eaton and Kardos 1972; Kircheis et al., 2004; Kircheis and Stanley 1981). In the Great Lakes, rainbow smelt declines in the mid-1900s were associated with alewife increases (Smith, 1968), also suggesting competition. However, more recent evaluations of alewife effects on rainbow smelt in the Great Lakes led to the conclusion that alewife are likely to have negligible impacts on rainbow smelt (Madenjian et al., 2008). Larger, deeper systems may reduce spatial overlap of alewife and rainbow smelt due to thermal structure while providing greater or more diverse zooplankton resources (Amsinck et al., 2006; Dodson, 1992; Simonin et al., 2012). In Lake Champlain, available oxythermal habitat was much smaller in the two smaller basins than in the Main Lake, where habitat was suitable at all depths and years. In the two smaller basins, warm epilimnetic waters and the hypoxic bottom layer during the warmest months (July to October) resulted in habitat constriction. Prior to the alewife invasion, rainbow smelt could reside in suboptimal warm water during summer habitat constriction without competition from alewife. Post-invasion occupancy of shallower waters during habitat constriction likely increased their overlap with alewife (Simonin et al., 2019). Additionally, *Mysis diluviana* which is a major diet item of rainbow smelt in Lake Champlain and the Great Lakes (Labar, 1993; Gamble et al., 2011a,b) and also consumed by alewife (Madenjian et al., 2003), is abundant in the Main Lake but virtually absent in the two smaller basins (Ball et al., 2015; Hrycik et al., 2015; O’Malley and Stockwell, 2019; JDS, pers. obs.). Consequently, *Mysis* may serve as a buffer to reduce competition between rainbow smelt and alewife in the Main Lake but not in the two smaller basins. In the absence of major changes in zooplankton density and body size, competition for zooplankton does not explain the declines in the smaller basins.

System size and habitat availability may also interact with predation to negatively influence rainbow smelt following alewife invasion and contribute to the patterns we observed. Larger systems may provide greater temporal and spatial mis-match between spawning adult alewife moving inshore and post-hatch larval rainbow smelt dispersal into large volumes of water offshore (dilution effect) and also provide stronger thermal gradients that promote vertical segregation (Madenjian et al., 2008; Recksiek and McCleave, 1973; Simonin et al., 2019). However, in the absence of evidence that alewife prey on larval and YOY rainbow smelt, the effect of scale may only be relevant to cannibalism and predation by large piscivores. Cannibalism could contribute to the apparent increased mortality of age-0 and −1 rainbow smelt we observed in the Northeast Arm and Malletts Bay. Cannibalism was in fact higher in Malletts Bay and the Northeast Arm than the Main Lake prior to the alewife invasion (Stetter Parker et al., 2007), but this did not appear to negatively affect abundance compared with the Main Lake. Cannibalism could only explain the decrease in rainbow smelt abundance after alewife invasion if increased competition with alewife forced rainbow smelt to increase cannibalism. Therefore, habitat scale and physicochemical constraints may have indirectly contributed to driving population declines in the two smaller basins if individuals were forced into habitats with more predators or competitors in the two smaller basins (Costantini et al., 2008; Horppila et al., 2003, 2004).

Other changes in the Lake Champlain ecosystem may have influenced rainbow smelt or influenced the effects of alewife. Portions of the lake have become more eutrophic over the past few decades, but only in shallow bays not suitable for rainbow smelt (Smeltzer et al. 2012). Of the 51 species that have invaded the lake, most do not overlap ecologically or geographically with rainbow smelt; e.g., invasive macrophytes are confined to the southern lake and littoral waters. Two possible exceptions are white perch (*Morone americana*) and zebra mussels (*Dreissena polymorpha*) that invaded the southern lake in 1984 and 1993, respectively, and spread rapidly throughout the Main Lake (Marsden and Hauser 2009). White perch are omnivorous and consume fish (Couture and Watzin, 2008; Schaeffer and Margraf, 1986) and therefore represent a predation threat. However, we should have observed demographic changes prior to the alewife invasion if white perch had a negative impact on rainbow smelt. Zebra mussel filtration lowers phytoplankton density and increases water transparency, leading to zooplankton declines (MacIsaac, 1996); however, Secchi disk readings in Lake Champlain increased only slightly and only in the south lake (Smeltzer et al., 2012) and adult densities remain too low in Malletts Bay and the Northeast Arm to expect them to have an impact (Marsden et al., 2013; VTDEC, 2020). No other changes to the lake have occurred with a timeline and magnitude that are likely to explain the changes we observed in rainbow smelt populations.

The successful invasion and rapid expansion of alewife in Lake Champlain is surprising, given the relatively intact fish community and high predator abundance. In the Great Lakes, alewife expanded soon after large piscivore and plantivore populations collapsed (Applegate and Van Meter, 1970; Baldwin et al., 2009; Miller, 1957; Smith, 1970). Alewife populations subsequently declined following sustained stocking of salmonines in the 1960s (Stewart and Ibarra, 1991). However, salmonines have been stocked continuously in Lake Champlain since the 1970s and at higher densities than in the Great Lakes. Even after reductions in lake trout stocking in 1995, lake trout plus Atlantic salmon stocking densities were 3.6 - 6 times higher per unit volume in Lake Champlain than in lakes Michigan or Huron (Great Lakes Fish Stocking database, www.glfc.org/fishstocking/; Stewart and Ibarra, 1991; Wehse et al., 2017). Both species began to consume alewife at least by 2008, when anglers and state biologists noted alewife in salmonine stomachs (Pientka, unpublished observations). By 2011, stable isotope analysis confirmed that alewife were a major element in Atlantic salmon and walleye diets and to a lesser extent in lake trout diets (Simonin et al., 2018). The presence of a robust rainbow smelt population in Lake Champlain would also be expected to potentially suppress the invasion, as rainbow smelt are predators of age-0 and yearling alewife (Foltz and Norden 1977; O’Gorman, 1974). Alternatively, the relatively simple planktivore community may have had low invasion resistance and provided a resource opportunity for alewife (Shea et al., 2002).

Neither zebra mussels nor predator abundance and food availability appear to be directly responsible for the rainbow smelt decline in the Northeast Arm and Malletts Bay. However, resource competition between rainbow smelt and alewife during summer may have been mediated by limitations in oxythermal habitat availability and/or alternative food resources (i.e., *Mysis*). Lake Champlain’s food web has been further modified since the forage fish survey was terminated in 2015. Sustained recruitment of wild lake trout was observed starting in 2015, and increased from 28% of the age-0 to age-3 population in 2015 to 66% in 2018 (Wilkins and Marsden in revision), increasing the abundance of top predators in the lake as stocking rates have remained constant. Juvenile lake trout target small rainbow smelt and alewife by age-1 (Marsden, Marden, unpublished data). Two new predatory cladocerans invaded the lake in 2014 (*Bythotrephes longimanus*) and 2018 (*Cercopagis pengoi*). Both invertebrate invaders compete with and are preyed on by rainbow smelt and alewife (Kozuchowski, 1996; Parker Stetter et al., 2005; Pothoven and Vanderploeg, 2004). Quagga mussel (*Dreissena bugensis*) invasion into Lake Champlain is likely, and this species could replace zebra mussels as they have in the Great Lakes (Mills et al 1999; Nalepa et al., 2010). While zebra mussels do not appear to have had an impact on rainbow smelt in Lake Champlain, quagga mussels have higher filtration rates and colonize deeper areas. Combined, all of these changes are likely to impact rainbow smelt and the Lake Champlain ecosystem in complex ways (e.g., Bunnell et al., 2012; Goto et al., 2020; Johannsson et al., 2011). Food web models and food web studies, in tandem, that compare basins will help better elucidate the potential impacts and pathways of system changes on Lake Champlain fish populations.

To summarize, we investigated the specific situation where rainbow smelt is native and alewife invasive. The native/introduced status of the two species was not a predictor of the impact of alewife on rainbow smelt. Instead, our results emphasize that the impact of alewife is context-dependent. Of the potential mechanisms to explain why rainbow smelt responded differently to an alewife invasion in the Main Lake than in the smaller basins, habitat size is the most convincing. Habitat size is known to be an important factor in the success or failure of a species to invade a system (Tamayo and Olden, 2014; Vander Zanden et al., 2004). Our findings indicate that habitat size may also play in an important role in the relative impact of invasive species. Consequently, managers must not only think about which systems are more vulnerable to invasion, but also which systems, once invaded, are the most likely to be impacted by the addition of invasive species.

## Acknowledgements

We thank George W. Labar who initiated the forage fish survey in the 1980s, and the captains and crews of the research vessels Melosira and Doré who conducted the surveys from 1984 to 1998 and from 1998 to 2015, respectively. We thank Angela Shambaugh and Pete Stangel at the VTDEC for providing us with the multiprobe data from the Lake Champlain long-term water quality and biological monitoring project. The project itself is supported by VTDEC, the NYDEC, and the Lake Champlain Basin Program. We thank the Marsden and Stockwell labs for their comments on an earlier draft of this manuscript. This project was made possible by fishing license sales and matching Dingell-Johnson/Wallop-Breaux funds, available through the Federal Aid in Sport Fish Restoration Act, and was supported by funding from Lake Champlain Sea Grant.

## Authors contribution

RB, JEM, JDS: Conceptualization; RB, BP, NS, TM: Data curation; RB, JEM, JDS: Formal analysis; RB, JEM, JDS: Funding acquisition; RB, JEM, JDS: Methodology; RB: Visualization. RB, JEM and JDS wrote the original draft; all authors reviewed, edited, and approved the final version of the manuscript.

## References

Amsinck, S.L., Strzelczak, A., Bjerring, R., Landkildehus, F., Lauridsen, T.L., Christoffersen, K., Jeppesen, E., 2006. Lake depth rather than fish planktivory determines cladoceran community structure in Faroese lakes – evidence from contemporary data and sediments. Freshw. Biol. 51, 2124–2142.

Applegate, V.C., Van Meter, H.D., 1970. A brief history of commercial fishing in Lake Erie (Federal Government Series No. 630), Fishery Leaflet. U.S. Fish and Wildlife Service, Washington, DC.

Baldwin, N.S., Saalfeld, R.W., Dochoda, M.R., Buettner, H.J., Eshenroder, R.L., 2009. Commercial fish production in the Great Lakes 1867-2006. [online]. Available from http://www.glfc.org/databases/commercial/commerc.php.

Ball, S.C., Mihuc, T.B., Myers, L.W., Stockwell, J.D., 2015. Ten-fold decline in *Mysis diluviana* in Lake Champlain between 1975 and 2012. J. Great Lakes Res. 41, 502–509.

Bigelow, H.B., Schroeder, W.C., 2002. Fishes of the Gulf of Maine: Fishery Bulletin 74. Blackburn Press.

Brandt, S.B., 1980. Spatial segregation of adult and young-of-the-year alewives across a thermocline in Lake Michigan. Trans. Am. Fish. Soc. 109, 469–478.

Brook, B., Sodhi, N., Bradshaw, C., 2008. Synergies among extinction drivers under global change. Trends Ecol. Evol. 23, 453–460.

Bunnell, D.B., Keeler, K.M., Puchala, E.A., Davis, B.M., Pothoven, S.A., 2012. Comparing seasonal dynamics of the Lake Huron zooplankton community between 1983–1984 and 2007 and revisiting the impact of *Bythotrephes* planktivory. J. Great Lakes Res. 38, 451–462.

Costantini, M., Ludsin, S.A., Mason, D.M., Zhang, X., Boicourt, W.C., Brandt, S.B., 2008. Effect of hypoxia on habitat quality of striped bass (*Morone saxatilis*) in Chesapeake Bay. Can. J. Fish. Aquat. Sci. 65, 989–1002.

Couture, S.C., Watzin, M.C., 2008. Diet of invasive adult white perch (*Morone americana*) and their effects on the zooplankton community in Missisquoi Bay, Lake Champlain. J. Great Lakes Res. 34, 485–494.

Dodson, S., 1992. Predicting crustacean zooplankton species richness. Limnol. Oceanogr. 37, 848–856.

Eaton, S.W., Kardos, L.P., 1971. The fishes of Canandaigua Lake. Sci. Stud. 28, 23–29.

Euclide, P.T., Pientka, B., Marsden, J. E., 2020. Genetic versus demographic stock structure of rainbow smelt in a large fragmented lake. J. Great Lakes Res.

Evans, D.O., Loftus, D.H., 1987. Colonization of inland lakes in the Great Lakes region by rainbow smelt, *Osmerus mordax*: their freshwater niche and effects on indigenous fishes. Can. J. Fish. Aquat. Sci. 44, s249–s266.

Fisheries Technical Committee, 2016. 2015 Annual Report. Lake Champlain Fish and Wildlife Management Cooperative.

Fisheries Technical Committee, 2009. 2008 Annual Report. Lake Champlain Fish and Wildlife Management Cooperative.

Foltz, J.W., Norden, C.R., 1977. Food habits and feeding chronology of rainbow smelt (*Osmerus mordax*) in Lake Michigan. Fish. Bull. 75, 637–640.

Fraga, S., Rodriguez, F., Bravo, I., Zapata, M., Marañon, E., 2012. Review of the main ecological features affecting benthic dinoflagellate blooms. Cryptogamie, Algologie 33, 171–179.

Glon, M.G., Larson, E.R., Reisinger, L.S., Pangle, K.L., 2017. Invasive dreissenid mussels benefit invasive crayfish but not native crayfish in the Laurentian Great Lakes. J. Great Lakes Res. 43, 289–297.

Goto, D., Dunlop, E.S., Young, J.D., Jackson, D.A., 2020. Shifting trophic control of fishery–ecosystem dynamics following biological invasions. Ecological Applications.

Guy, C.S., Brown, M.L. (Eds.), 2007. Analysis and Interpretation of Freshwater Fisheries Data. American Fisheries Society. American Fisheries Society, Bethesda, Md.

Happel, A., Jonas, J.L., McKenna, P.R., Rinchard, J., He, J.X., Czesny, S.J., 2018. Spatial variability of lake trout diets in Lakes Huron and Michigan revealed by stomach content and fatty acid profiles. Can. J. Fish. Aquat. Sci. 75, 95–106.

Harder, A.M., Ardren, W.R., Evans, A.N., Futia, M.H., Kraft, C.E., Marsden, J.E., Richter, C.A., Rinchard, J., Tillitt, D.E., Christie, M.R., 2018. Thiamine deficiency in fishes: causes, consequences, and potential solutions. Rev. Fish Biol. Fisheries 28, 865–886.

He, X., Labar, G.W., 1994. Interactive effects of cannibalism, recruitment, and predation on rainbow smelt in Lake Champlain: a modeling synthesis. J. Great Lakes Res. 20, 289–298.

Horppila, J., Liljendahl-Nurminen, A., Malinen, T., 2004. Effects of clay turbidity and light on the predatorprey interaction between smelts and chaoborids. Can. J. Fish. Aquat. Sci. 61, 1862–1870.

Horppila, J., Liljendahl-Nurminen, A., Malinen, T., Salonen, M., Tuomaala, A., Uusitalo, L., Vinni, M., 2003. *Mysis relicta* in a eutrophic lake: Consequences of obligatory habitat shifts. Limnol. Oceanogr. 48, 1214–1222.

Hrycik, A.R., Almeida, L.Z., Höök, T.O., 2017. Sub-lethal effects on fish provide insight into a biologically-relevant threshold of hypoxia. Oikos 126, 307–317.

Hrycik, A.R., Simonin, P.W., Rudstam, L.G., Parrish, D.L., Pientka, B., Mihuc, T.B., 2015. Mysis zooplanktivory in Lake Champlain: A bioenergetics analysis. J. Great Lakes Res. 41, 492–501.

Jacobs, G.R., Bruestle, E.L., Hussey, A., Gorsky, D., Fisk, A.T., 2017. Invasive species alter ontogenetic shifts in the trophic ecology of Lake Sturgeon (*Acipenser fulvescens*) in the Niagara River and Lake Ontario. Biol. Invasions 19, 1533–1546.

Johannsson, O.E., Bowen, K.L., Holeck, K.T., Walsh, M.G., 2011. *Mysis diluviana* population and cohort dynamics in Lake Ontario before and after the establishment of *Dreissena* spp., *Cercopagis pengoi*, and *Bythotrephes longimanus*. Can. J. Fish. Aquat. Sci. 68, 795–811.

Kircheis, F.W., Stanley, J.G., 1981. Theory and practice of forage-fish management in New England. Trans. Am. Fish. Soc. 110, 729–737.

Kircheis, F.W., Trial, J.G., Boucher, D.P., Mower, B., Squiers, T., Gray, N., O’Donnel, M., Stahlnecker, J., 2004. Analysis of impacts related to the introduction of anadromous alewives into a small freshwater lake in central Maine, USA. Maine Department of Inland Fisheries and Wildlife, Bangor, Maine.

Kirn, R.A., Labar, G.W., 1996. Growth and survival of rainbow smelt, and their role as prey for stocked salmonids in Lake Champlain. Trans. Am. Fish. Soc. 125, 87–96.

Kozuchowski, E.S., 1996. Prey selection by rainbow smelt in eastern Lake Erie following the introduction of *Bythotrephes cederstroemi* (Master Thesis). State University of New York, College at Buffalo, Buffalo, NY.

Krueger, C.C., Perkins, D.L., Mills, E.L., Marsden, J. E., 1995. Predation by alewives on lake trout fry in Lake Ontario: Role of an exotic species in preventing restoration of a native species. J. Great Lakes Res., 21 (suppl. 1), 458–469.

Labar, G.W., 1998. Assessment of rainbow smelt stocks during an eight-year experimental sea lamprey control program on Lake Champlain. Vermont Department of Fish and Wildlife, Essex Junction, VT.

Labar, G.W., 1993. Use of bioenergetics models to predict the effect of increased lake trout predation on rainbow smelt following sea lamprey control. Trans. Am. Fish. Soc. 122, 942–950.

Lake Champlain Basin Program, 2018. State of the lake and ecosystem indicators report. Lake Champlain Basin Program, Grand Isle, VT

Latta, W.C., 1995. Distribution and abundance of lake herring (*Coregonus artedi*) in Michigan. (Fisheries research report No. 2014). Michigan Department of Natural Resources, Fisheries Division, Ann Arbor, MI.

Lantry, B.F., Stewart, D.J., 1993. Ecological energetics of rainbow smelt in the Laurentian Great Lakes: an interlake comparison. Transactions of the American Fisheries Society 122, 951–976.

MacIsaac, H.J., 1996. Potential abiotic and biotic impacts of zebra mussels on the inland waters of North America. Am. Zool. 36, 287–299.

Mackie, G.L., Schloesser, D.W., 1996. Comparative biology of zebra mussels in Europe and North America: an overview. Am. Zool. 36:3, 244–258.

Madenjian, C.P., Holuszko, J.D., Desorcie, T.J., 2003. Growth and condition of alewives in Lake Michigan, 1984–2001. Trans. Am. Fish. Soc. 132, 1104–1116.

Madenjian, C.P., O’Gorman, R., Bunnell, D.B., Argyle, R.L., Roseman, E.F., Warner, D.M., Stockwell, J.D., Stapanian, M.A., 2008. Adverse effects of alewives on Laurentian Great Lakes Fish communities. N. Am. J. Fish. Manag. 28, 263–282.

Marsden, J.E., Chipman, B.D., Nashett, L.J., Anderson, J.K., Bouffard, W., Durfey, L., Gersmehl, J.E., Schoch, W.F., Staats, N.R., Zerrenner, A., 2003. Sea Lamprey control in Lake Champlain. J. Great Lakes Res. 29, 655–676.

Marsden, J.E., Hauser, M., 2009. Exotic species in Lake Champlain. J. Great Lakes Res. 35, 250–265.

Marsden, J.E., Langdon, R.W., 2012. The history and future of Lake Champlain’s fishes and fisheries. J. Great Lakes Res. 38, 19–34.

Marsden, J.E., Stangel, P., Shambaugh, A., 2013. Influence of environmental factors on zebra mussel population expansion in Lake Champlain, 1994–2010, in: Quagga and Zebra Mussels. CRC Press, pp. 33–54.

Mihuc, T.B., Dunlap, F., Binggeli, C., Myers, L., Pershyn, C., Groves, A., Waring, A., 2012. Long-term patterns in Lake Champlain’s zooplankton: 1992–2010. J. Great Lakes Res. 38, 49–57.

Miller, R.R., 1957. Origin and dispersal of the alewife, *Alosa pseudoharengus*, and the gizzard shad, *Dorosoma cepedianum*, in the Great Lakes. Trans. Am. Fish. Soc. 86, 97–111.

Mills, E.L., Leach, J.H., Carlton, J.T., Secor, C.L., 1993. Exotic species in the Great Lakes: a history of biotic crises and anthropogenic introductions. J. Great Lakes Res. 19, 1–54.

Myers, J.T., Jones, M.L., Stockwell, J.D., Yule, D.L., 2009. Reassessment of the predatory effects of rainbow smelt on ciscoes in Lake Superior. Trans. Am. Fish. Soc. 138, 1352–1368.

Ogle, D., Wheeler, P., Dinno, A., 2020. FSA: simple fisheries stock assessment methods.

O’Gorman, R., 1974. Predation by rainbow smelt (*Osmerus mordax*) on young-of-the-year alewives (*Alosa pseudoharengus*) in the Great Lakes. Prog. Fish. Cult. 36, 223–224.

O’Malley, B.P., Stockwell, J.D., 2019. Diel feeding behavior in a partially migrant *Mysis* population: A benthic-pelagic comparison. Food Webs e00117.

Parker Stetter, S.L., Stritzel Thomson, J.L., Rudstam, L.G., Parrish, D.L., Sullivan, P.J., 2007. Importance and predictability of cannibalism in rainbow smelt. Trans. Am. Fish. Soc. 136, 227–237.

Parker Stetter, S.L., Witzel, L.D., Rudstam, L.G., Einhouse, D.W., Mills, E.L., 2005. Energetic consequences of diet shifts in Lake Erie rainbow smelt (*Osmerus mordax*). Can. J. Fish. Aquat. Sci. 62, 145–152.

Peterson, B.G., Carl, P., 2020. PerformanceAnalytics: Econometric tools for performance and risk analysis. R package version 2.0.4.

Pothoven, S.A., Vanderploeg, H.A., 2004. Diet and prey selection of alewives in Lake Michigan: seasonal, depth, and interannual patterns. Trans. Am. Fish. Soc. 133, 1068–1077.

Power, M.E., 1992. Top-down and bottom-up forces in food webs: do plants have primacy. Ecology 73, 733–746.

R Core Team, 2020. R: A language and environment for statistical computing. R Foundation for Statistical Computing, Vienna, Austria. R Foundation for Statistical Computing.

Ray, B.A., Hrabik, T.R., Ebener, M.P., Gorman, O.T., Schreiner, D.R., Schram, S.T., Sitar, S.P., Mattes, W.P., Bronte, C.R., 2007. Diet and prey selection by Lake Superior lake trout during spring, 1986– 2001. J. Great Lakes Res. 33, 104–113.

Recksiek, C.W., McCleave, J.D., 1973. Distribution of pelagic fishes in the Sheepscot River - Black River estuary. Trans. Am. Fish. Soc. 102, 541–551.

Ricciardi, A., MacIsaac, H.J., 2010. Impacts of biological invasions on freshwater ecosystems, in: Fifty Years of Invasion Ecology. John Wiley & Sons, Ltd, pp. 211–224.

Ricciardi, A. 2006. Patterns of invasion in the Laurentian Great Lakes in relation to changes in vector activity. Divers. Distrib. 12, 425–433.

Ricker, W.E., 1975. Computation and interpretation of biological statistics of fish populations. Bulletin of the Fisheries Research Board of Canada 191, 1–382.

Schaeffer, J.S., Margraf, F.J., 1987. Predation on fish eggs by white perch, *Morone americana*, in western Lake Erie. Environ. Biol. Fish. 18, 77–80.

Simberloff, D., Von Holle, B., 1999. Positive interactions of nonindigenous species: invasional meltdown? Biol. Invasions 1, 21–32.

Simonin, P.W., Parrish, D.L., Rudstam, L.G., Sullivan, P.J., Pientka, B., 2012. Native rainbow smelt and nonnative alewife distribution related to temperature and light gradients in Lake Champlain. J. Great Lakes Res., Lake Champlain in 2010. 38, 115–122.

Simonin, P.W., Rudstam, L.G., Parrish, D.L., Pientka, B., Sullivan, P.J., 2018. Piscivore diet shifts and trophic level change after alewife establishment in Lake Champlain. Trans. Am. Fish. Soc. 147, 939–947.

Simonin, P.W., Rudstam, L.G., Sullivan, P.J., Parrish, D.L., Pientka, B., 2019. Early mortality and freshwater forage fish recruitment: nonnative alewife and native rainbow smelt interactions in Lake Champlain. Can. J. Fish. Aquat. Sci. 76, 806–814.

Smeltzer, E., Shambaugh, A. d., Stangel, P., 2012. Environmental change in Lake Champlain revealed by long-term monitoring. J. Great Lakes Res. 38, 6–18.

Smith, S.H., 1970. Species interactions of the alewife in the Great Lakes. Trans. Am. Fish. Soc. 99, 754–765.

Smith, S.H., 1968. Species succession and fishery exploitation in the Great Lakes. J. Fish. Res. Bd. Can. 25:4, 667–693.

Stewart, D.J., Binkowski, F.P., 1986. Dynamics of consumption and food conversion by Lake Michigan alewives: an energetics-modeling synthesis. Trans. Am. Fish. Soc. 115, 643–661.

Stewart, D.J., Ibarra, M., 1991. Predation and production by salmonine fishes in Lake Michigan, 1978–88. Can. J. Fish. Aquat. Sci. 48, 909–922.

Stewart, D.J., Kitchell, J.F., Crowder, L.B., 1981. Forage fishes and their salmonid predators in Lake Michigan. Trans. Am. Fish. Soc. 110:6, 751–763.

Stewart, T.J., Sprules, W.G., O’Gorman, R., 2009. Shifts in the diet of Lake Ontario alewife in response to ecosystem change. J. Great Lakes Res. 35, 241–249.

Stritzel Thomson, J.L., Parrish, D.L., Parker-Stetter, S.L., Rudstam, L.G., Sullivan, P.J., 2011. Growth rates of rainbow smelt in Lake Champlain: effects of density and diet: Growth rates of rainbow smelt. Ecol. Freshw. Fish 20, 503–512.

Tamayo, M., Olden, J.D., 2014. Forecasting the Vulnerability of Lakes to Aquatic Plant Invasions. Invas. Plant Sci. Mana. 7, 32–45.

Urban, T.P., Brandt, S.B., 1993. Food and habitat partitioning between young-of-year alewives and rainbow smelt in southeastern Lake Ontario. Environ. Biol. Fish. 36, 359–372.

VTDEC, 2020. Lake Champlain long-term monitoring project. Vermont Department of Environmental Conservation, Montpellier, VT.

Wehse, R., Hanson, D., Treska, T., Holey, M., 2017. Summary of 2016 lake trout and salmonid stocking in Lake Michigan (No. 2017– 02). U.S. Fish and Wildlife Service. Green Bay Fish and Wildlife Conservation Office, New Franken, WI.

Wickham, H., 2016. ggplot2: elegant graphics for data analysis, Second edition. ed, Use R! Springer-Verlag, New York.

