## Appendix A for "Rainbow smelt population responses to species invasions and change in environmental condition"

### Appendix A — Results of Kruskal-Wallis pairwise comparison test.

**Table S1.** Results of Kruskal-Wallis pairwise comparison test between periods for the three basins and the four variables of interest. *p* is the overall p-value, *p.adj* is the p-value with Bonferroni correction to account for multiple comparison. *p.signif* indicates the level of significance: ns (p.adj > 0.05); * (p.adj > 0.01); ** (p.adj > 0.001); *** (p.adj > 0.0001); **** (p.adj ≤ 0.0001).

| variable | Basin | Period 1 | Period 2 | p | p.adj | p.signif |
|---|---|---|---|---|---|---|
| log(CPUE) | Main Lake | 1987-2002 | 2003-2006 | 0.362 | 1.000 | ns |
| | | 1987-2002 | 2007-2015 | 0.688 | 1.000 | ns |
| | | 2003-2006 | 2007-2015 | 0.727 | 1.000 | ns |
| | Malletts Bay | 1987-2002 | 2003-2006 | 0.018 | 0.053 | ns |
| | | 1987-2002 | 2007-2015 | 0.000 | 0.000 | **** |
| | | 2003-2006 | 2007-2015 | 0.003 | 0.008 | ** |
| | Northeast Arm | 1987-2002 | 2003-2006 | 0.442 | 1.000 | ns |
| | | 1987-2002 | 2007-2015 | 0.000 | 0.001 | *** |
| | | 2003-2006 | 2007-2015 | 0.008 | 0.024 | * |
| Average length age-2+ (mm) | Main Lake | 1987-2002 | 2003-2006 | 0.412 | 1.000 | ns |
| | | 1987-2002 | 2007-2015 | 0.209 | 0.630 | ns |
| | | 2003-2006 | 2007-2015 | 1.000 | 1.000 | ns |
| | Malletts Bay | 1987-2002 | 2003-2006 | 0.130 | 0.390 | ns |
| | | 1987-2002 | 2007-2015 | 0.324 | 0.970 | ns |
| | | 2003-2006 | 2007-2015 | 0.050 | 0.150 | ns |
| | Northeast Arm | 1987-2002 | 2003-2006 | 0.060 | 0.180 | ns |
| | | 1987-2002 | 2007-2015 | 0.000 | 0.000 | *** |
| | | 2003-2006 | 2007-2015 | 0.034 | 0.100 | ns |
| Average condition (age 2-3) | Main Lake | 1987-2002 | 2003-2006 | 0.549 | 1.000 | ns |
| | | 1987-2002 | 2007-2015 | 0.896 | 1.000 | ns |
| | | 2003-2006 | 2007-2015 | 1.000 | 1.000 | ns |

| | | | | | | |
|---|---|---|---|---|---|---|
| | Malletts Bay | 1987-2002 | 2003-2006 | 0.477 | 1.000 | ns |
| | | 1987-2002 | 2007-2015 | 0.357 | 1.000 | ns |
| | | 2003-2006 | 2007-2015 | 0.260 | 0.780 | ns |
| | Northeast Arm | 1987-2002 | 2003-2006 | 0.871 | 1.000 | ns |
| | | 1987-2002 | 2007-2015 | 0.126 | 0.380 | ns |
| | | 2003-2006 | 2007-2015 | 0.503 | 1.000 | ns |
| Annual mortality | Main Lake | 1987-2002 | 2003-2006 | 0.839 | 1.000 | ns |
| | | 1987-2002 | 2007-2015 | 0.635 | 1.000 | ns |
| | | 2003-2006 | 2007-2015 | 0.762 | 1.000 | ns |
| | Malletts Bay | 1987-2002 | 2003-2006 | 0.002 | 0.007 | ** |
| | | 1987-2002 | 2007-2015 | 0.945 | 1.000 | ns |
| | | 2003-2006 | 2007-2015 | 0.114 | 0.340 | ns |
| | Northeast Arm | 1987-2002 | 2003-2006 | 0.102 | 0.310 | ns |
| | | 1987-2002 | 2007-2015 | 0.521 | 1.000 | ns |
| | | 2003-2006 | 2007-2015 | 0.257 | 0.770 | ns |

**Table S2.** Results of Kruskal-Wallis pairwise comparison test between periods for the three basins and the zooplankton taxa of interest. *p* is the overall p-value, *p.adj* is the p-value with Bonferroni correction. *p.signif* indicates the level of significance: ns, p>0.05; *, p>0.01; **, p>0.001; ***, p>0.0001; ****: p≤0.0001

| variable | Taxa | Basin | Period 1 | Period 2 | p | p.adj | p.signif |
|---|---|---|---|---|---|---|---|
| Mean summer density (#/ $m^3$) | Cyclopoida | Main Lake | 1992-2002 | 2003-2006 | 0.104 | 0.310 | ns |
| | | | 1992-2002 | 2007-2015 | 0.230 | 0.690 | ns |
| | | | 2003-2006 | 2007-2015 | 0.050 | 0.150 | ns |
| | | Malletts Bay | 1992-2002 | 2003-2006 | 0.476 | 1.000 | ns |
| | | | 1992-2002 | 2007-2015 | 0.067 | 0.200 | ns |
| | | | 2003-2006 | 2007-2015 | 0.020 | 0.059 | * |
| | | Northeast Arm | 1992-2002 | 2003-2006 | 0.661 | 1.000 | ns |
| | | | 1992-2002 | 2007-2015 | 0.976 | 1.000 | ns |
| | | | 2003-2006 | 2007-2015 | 0.446 | 1.000 | ns |
| | Calanoida | Main Lake | 1992-2002 | 2003-2006 | 0.001 | 0.004 | ** |
| | | | 1992-2002 | 2007-2015 | 0.056 | 0.170 | ns |
| | | | 2003-2006 | 2007-2015 | 0.003 | 0.008 | ** |
| | | Malletts Bay | 1992-2002 | 2003-2006 | 0.762 | 1.000 | ns |
| | | | 1992-2002 | 2007-2015 | 0.616 | 1.000 | ns |
| | | | 2003-2006 | 2007-2015 | 0.446 | 1.000 | ns |
| | | Northeast Arm | 1992-2002 | 2003-2006 | 0.056 | 0.170 | ns |
| | | | 1992-2002 | 2007-2015 | 0.413 | 1.000 | ns |
| | | | 2003-2006 | 2007-2015 | 0.030 | 0.089 | ns |
| | Bosminidae | Main Lake | 1992-2002 | 2003-2006 | 0.851 | 1.000 | ns |
| | | | 1992-2002 | 2007-2015 | 0.295 | 0.880 | ns |
| | | | 2003-2006 | 2007-2015 | 0.825 | 1.000 | ns |
| | | Malletts Bay | 1992-2002 | 2003-2006 | 0.610 | 1.000 | ns |
| | | | 1992-2002 | 2007-2015 | 0.682 | 1.000 | ns |
| | | | 2003-2006 | 2007-2015 | 1.000 | 1.000 | ns |
| | | Northeast Arm | 1992-2002 | 2003-2006 | 0.753 | 1.000 | ns |
| | | | 1992-2002 | 2007-2015 | 0.740 | 1.000 | ns |
| | | | 2003-2006 | 2007-2015 | 0.446 | 1.000 | ns |
| | Daphnidae | Main Lake | 1992-2002 | 2003-2006 | 0.003 | 0.009 | ** |
| | | | 1992-2002 | 2007-2015 | 0.412 | 1.000 | ns |
| | | | 2003-2006 | 2007-2015 | 0.003 | 0.008 | ** |
| | | Malletts Bay | 1992-2002 | 2003-2006 | 1.000 | 1.000 | ns |
| | | | 1992-2002 | 2007-2015 | 0.102 | 0.310 | ns |
| | | | 2003-2006 | 2007-2015 | 0.030 | 0.089 | ns |
| | | Northeast Arm | 1992-2002 | 2003-2006 | 0.489 | 1.000 | ns |
| | | | 1992-2002 | 2007-2015 | 0.566 | 1.000 | ns |
| | | | 2003-2006 | 2007-2015 | 0.862 | 1.000 | ns |
| | Rotifera | Main Lake | 1992-2002 | 2003-2006 | 0.343 | 1.000 | ns |

| | | | | | | | |
|---|---|---|---|---|---|---|---|
| | | | 1992-2002 | 2007-2015 | 0.080 | 0.240 | ns |
| | | | 2003-2006 | 2007-2015 | 0.604 | 1.000 | ns |
| | Mysids | Main Lake | 1992-2002 | 2003-2006 | 0.280 | 0.840 | ns |
| | | | 1992-2002 | 2007-2015 | 0.020 | 0.060 | ns |
| | | | 2003-2006 | 2007-2015 | 0.503 | 1.000 | ns |
| Mean length (mm) | *Diacyclops thomasi* | Main Lake | 2001-2002 | 2003-2006 | 0.267 | 0.800 | ns |
| | | | 2001-2002 | 2007-2015 | 0.288 | 0.860 | ns |
| | | | 2003-2006 | 2007-2015 | 0.588 | 1.000 | ns |
| | | Northeast Arm | 2001-2002 | 2003-2006 | 1.000 | 1.000 | ns |
| | | | 2001-2002 | 2007-2015 | 0.178 | 0.530 | ns |
| | | | 2003-2006 | 2007-2015 | 0.085 | 0.250 | ns |
| | *Daphnia retrocurva* | Main Lake | 2001-2002 | 2003-2006 | 0.533 | 1.000 | ns |
| | | | 2001-2002 | 2007-2015 | 0.045 | 0.130 | ns |
| | | | 2003-2006 | 2007-2015 | 0.017 | 0.050 | * |
| | | Northeast Arm | 2001-2002 | 2003-2006 | 0.400 | 1.000 | ns |
| | | | 2001-2002 | 2007-2015 | 0.694 | 1.000 | ns |
| | | | 2003-2006 | 2007-2015 | 0.184 | 0.550 | ns |
| | *Leptodiaptomus sicilis* | Main Lake | 2001-2002 | 2003-2006 | 0.800 | 1.000 | ns |
| | | | 2001-2002 | 2007-2015 | 0.723 | 1.000 | ns |
| | | | 2003-2006 | 2007-2015 | 0.315 | 0.950 | ns |
| | | Northeast Arm | 2001-2002 | 2003-2006 | 0.236 | 0.710 | ns |
| | | | 2001-2002 | 2007-2015 | 0.190 | 0.570 | ns |
| | | | 2003-2006 | 2007-2015 | 1.000 | 1.000 | ns |
| | *Bosmina longirostris* | Main Lake | 2001-2002 | 2003-2006 | 0.348 | 1.000 | ns |
| | | | 2001-2002 | 2007-2015 | 0.532 | 1.000 | ns |
| | | | 2003-2006 | 2007-2015 | 0.201 | 0.600 | ns |
| | | Northeast Arm | 2001-2002 | 2003-2006 | 0.800 | 1.000 | ns |
| | | | 2001-2002 | 2007-2015 | 0.507 | 1.000 | ns |
| | | | 2003-2006 | 2007-2015 | 0.304 | 0.910 | ns |
