## Appendix B for "Rainbow smelt population responses to species invasions and change in environmental condition"

### Appendix B — Temperature and DO profile

We used vertical profile data obtained with a multiprobes sensor by the Vermont Department of Environmental Conservation (DEC), as part of the Lake Champlain long-term monitoring program initiated in 1992 (https://dec.vermont.gov/watershed/lakes-ponds/monitor/lake-champlain). The program is carried on jointly with the New York DEC with fundings from Lake Champlain Basin Program and the two states. Fifteen stations are sampled fortnightly from late April to early November, and we selected stations 19, 25 and 34 to represent conditions in the Main Lake, Malletts Bay, and the Northeast Arm respectively (Fig. 1 main text).

We rounded the depths to the closest meter and built heatmaps using the function *filled.contour()* in R (Fig. S2-1, S2-2) (Read et al., 2011).

### Reference

Read, J.S., Hamilton, D.P., Jones, I.D., Muraoka, K., Winslow, L.A., Kroiss, R., Wu, C.H., Gaiser, E., 2011. Derivation of lake mixing and stratification indices from high-resolution lake buoy data. Environ. Model. Softw. 26, 1325–1336.

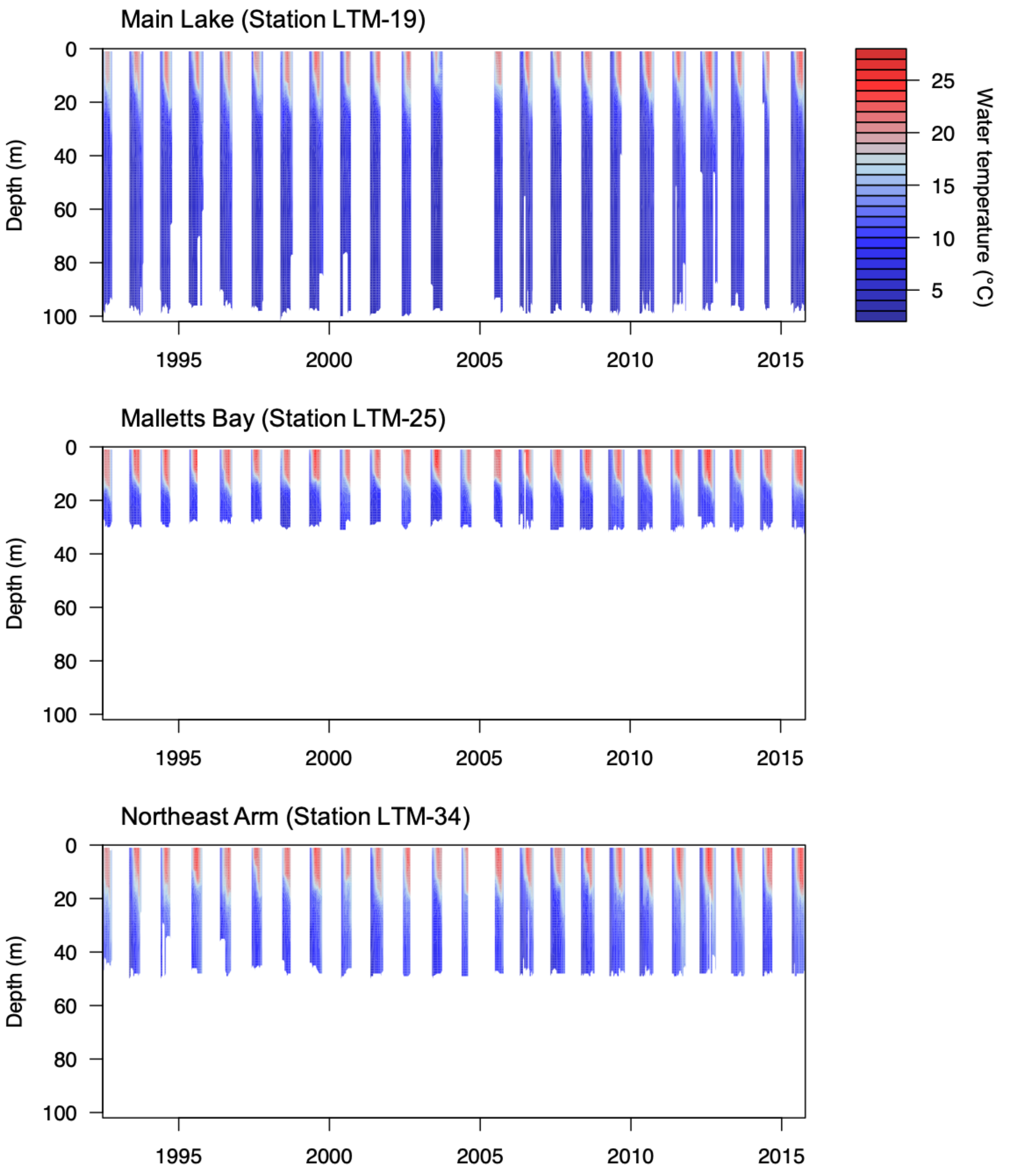


**Figure S2-1.** Water temperature (°C) in three basins of Lake Champlain each summer between 1992 and 2015.

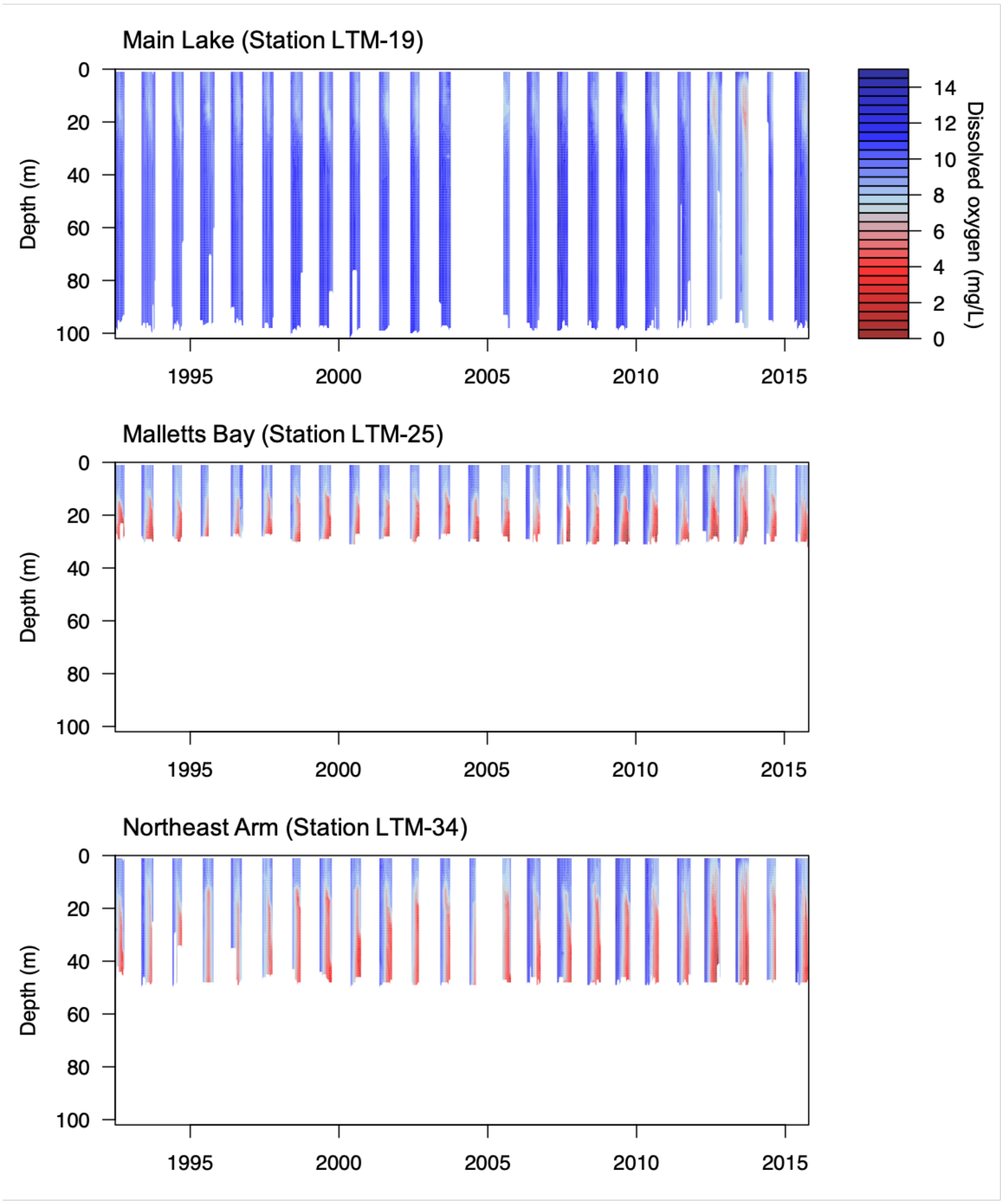


**Figure S2-2.** Dissolved oxygen (mg/l) in three basins of Lake Champlain each summer between 1992 and 2015.
